# Iran’s roads: mitigating the most serious threat to the critically endangered Asiatic cheetah *Acinonyx jubatus venaticus*

**DOI:** 10.1101/230581

**Authors:** Jamshid Parchizadeh, Maria Gatta, Roberta Bencini, Ali Turk Qashqaei, Mohammad Ali Adibi, Samual T. Williams

## Abstract

Wildlife-vehicle collisions are an important cause of mortality for many species, and the number of collisions is expected to grow rapidly as the global road network quickly expands over the next few decades. Wildlife-vehicle collisions also have the potential to be extremely detrimental to small wildlife populations, such as the critically endangered Asiatic cheetah (*Acinonyx jubatus venaticus*), with only 43 individuals remaining in the wild. We assessed the spatial distribution of road mortalities between 2004 and 2016 to identify roadkill hotspots involving Asiatic cheetahs in Iran using network kernel density estimation. A total of sixteen cheetah fatalities due to wildlife-vehicle collisions were recorded, and we identified six road fragments as roadkill hotspots. Efforts to reduce wildlife-cheetah collisions should be targeted in the densest hotspots. We review the options available to achieve this, and we recommend a strategic shift away from the ineffective warning signage currently used, and instead suggest adopting an evidence-based approach focusing on installing wildlife crossing structures in conjunction with fencing in roadkill hotspots. These measures will help to enhance the conservation status of the Asiatic cheetah, as the current high level of mortality of Asiatic cheetahs on Iran’s roads could have potentially dramatic impacts on this critically endangered subspecies.

## Introduction

The rapid expansion of infrastructure has had severe impacts on the abundance of wildlife across the globe (Grilo et al. 2015). Roads can be particularly harmful, as they have both indirect effects such as providing people with access for hunting and further development, and direct effects such as inhibiting the movement of animals, and causing direct mortality through wildlife-vehicle collisions (Laurance et al. 2009; Collinson et al. 2015). The taxa that are at the greatest risk to threats associated with roads include amphibians, reptiles, and mammals, especially large carnivores such as cheetahs (*Acinonyx jubatus*), due to their large home ranges and small population sizes (Grilo et al. 2015; Ceia-Hasse et al. 2017; Durant et al. 2017; M. Huijser, pers. comm.). Collisions with vehicles are known to be a source of cheetah mortality in countries such as Tanzania and South Africa (Tawiri 2005; Collinson et al. 2015), but this is rarely quantified in relation to other threats (Gadd 2015). In Iran the critically endangered Asiatic cheetah (*A. j. venaticus*) is particularly susceptible to these threats due to its extremely small population size (Jowkar et al. 2008).

A total of only 43 free-roaming Asiatic cheetahs remain (Durant et al. 2017), all of which occur in central Iran in the provinces of Isfahan, Kerman, North Khorasan, Semnan, South Khorasan, and Yazd (Khaleghparast 2015). After an absence of almost 40 years, Asiatic cheetahs were recently observed in the Golestan and Razavi Khorasan provinces (Khaleghparast 2015). Collisions with vehicles pose the most serious threat to Asiatic cheetahs, with one to two Asiatic cheetahs killed by vehicles on Iran’s roads annually (Hunter et al. 2007; Farhadinia et al. 2017). Between 2004 and 2016 roadkill was the most important cause of Asiatic cheetah mortality (Ahmadi et al. 2017). There have been few studies, however, of the distribution of cheetah roadkills, although this would be incredibly useful to inform mitigation strategies.

We assessed the distribution of road mortalities involving Asiatic cheetahs to establish where these events occur most frequently, enabling us to identify roadkill hotspots at which mitigation measures should be targeted. We also briefly review potential strategies that could be implemented to reduce the threats posed by roads to cheetahs in Iran.

## Methods

### Study area

Iran has a total area 1,648,195 km^2^, and a quarter of the country is composed of arid and semi-arid deserts (Mansouri 2004; Sabziparvar 2008) (Fig. 1). Kerman, Semnan, and Yazd Provinces are located between 51°47′20″ and 61°09′47″ E and 26°00′40″ and 37°21′54″ N, and cover an area of 480,909 km^2^. The climate of the provinces is dry, with mean maximum daily temperature of up to 55 °C in summer, and mean minimum daily temperatures of down to -20 °C in winter. Altitude varies from 500 to 1,200 m, with precipitation ranging from 70 to 111 mm annually. The dominant plant species are *Artemisia siberi* and *Astragalus gossypius* (Heshmati 2007).

**Fig. 1.**
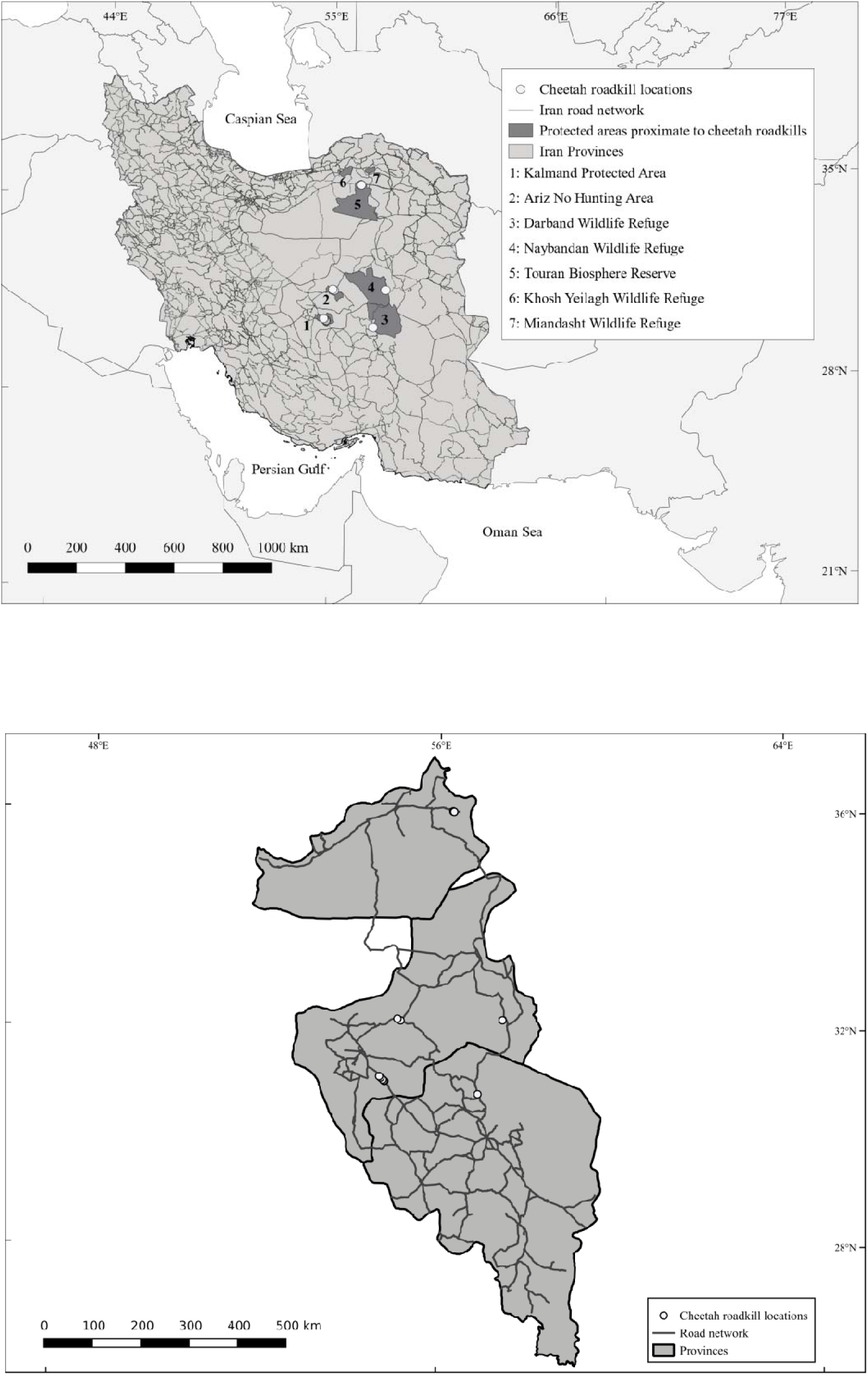
Map showing the locations of cheetah roadkill across a) Iran, and b) the three provinces in which the mortalities were recorded (the name of the provinces downward are; Semnan, Yazd, and Kerman) (some circles represent multiple cheetah roadkill locations)

### Data collection

Between 2004 and 2016 local people reported the location of cheetah carcasses they encountered along roads by telephoning their local Office of Iran’s Department of the Environment. The authorities inspected the carcasses to determine cause of death and recorded the coordinates using GPS Garmin Etrex 10 (Garmin International Inc., Olathe, KS, USA). The data were collected by the Conservation of the Asiatic Cheetah Project, a project under Iran’s Department of the Environment in collaboration with the United Nations Development Programme, the Wildlife Conservation Society, Panthera, Cheetah Conservation Fund, and the Cat Specialist Group of the International Union for Conservation of Nature (Conservation of the Asiatic Cheetah Project 2001).

### Data analysis

We used network kernel density estimation (NetKDE) to identify the locations of Asiatic cheetah roadkill hotspots, which is an extension of the planar KDE methods and has been frequently used to identify wildlife-vehicle collision hotspots for other species (Danks and Porter 2010; Morelle et al. 2013; Snow et al. 2014; Özcan and Ôzkazanç 2017). Planar KDE produces a probability surface of points in a 2-dimensional space, which provides a visualization of different concentrations of the event across the study area. A critical aspect of the KDE analysis is the bandwidth value (*h*) which is the distance over which a data point has influence. However, accidents along a road network do not meet the assumptions of homogeneity of 2-dimensional space (Xie and Yan 2008; Okabe et al. 2009). NetKDE has the advantage over planar KDE that distances are not measured in 2-dimensional Euclidean space, but rather in network distance. We performed these analyses using QGIS version 2.18.13 (Quantum GIS Development Team 2017) and SANET version 4.1 Standalone (Okabe et al. 2006).

Cheetah carcases were generally not located directly on road surfaces, so the closest point on the road network incorporated into the analyses. Most cheetah mortalities were located close to paved roads, but two were recorded near to unpaved roads, in which case the unpaved roads in close proximity to the data points were digitised using satellite imagery from Google Earth, and added to the road network. We performed the NetKDE analyses using the kernel type equal split at discontinuous nodes in SANET. Different bandwidths were tested and visually inspected (500 m, 750 m, 850 m, 1,000 m, 1,500 m, 2,000 m). We selected the 1,500 m bandwidth as the most suitable for our dataset, given the low sample size and large network area.

## Results

A total of 16 Asiatic cheetah roadkills were recorded between 2004 and 2016 in Iran (Fig. 1). Nine collisions with cheetahs occurred inside protected areas and 7 occurred on roads bordering protected areas. Most Asiatic cheetah road mortalities occurred in Yazd Province (9 roadkills), followed by Semnan Province (6 roadkills), and Kerman Province (1 roadkill; Table 1). The loss of females is particularly problematic, as this will exacerbate the population decline by reducing recruitment rates (Marker et al. 2003). A total of six road fragments were identified as hotspots for Asiatic cheetah road mortality. One road fragment hotspot contained six roadkills, another hotspot contained three roadkills, and four hotspots contained only one roadkill. Three cheetah road mortalities fell into areas that were not identified as roadkill hotspots. The road fragment made up of six cheetah collisions was the only hotspot located in Semnan Province. Five hotspots were located in Yazd Province, while no hotspots were located in Kerman Province (Fig. 2). The length of the road fragments identified as hotspots varied between 8 m and 3,863 m. The total length of the road networks that were identified as hotspots was 7.9 km, which represents 0.009% of the total road network analysed.

**Table 1.**
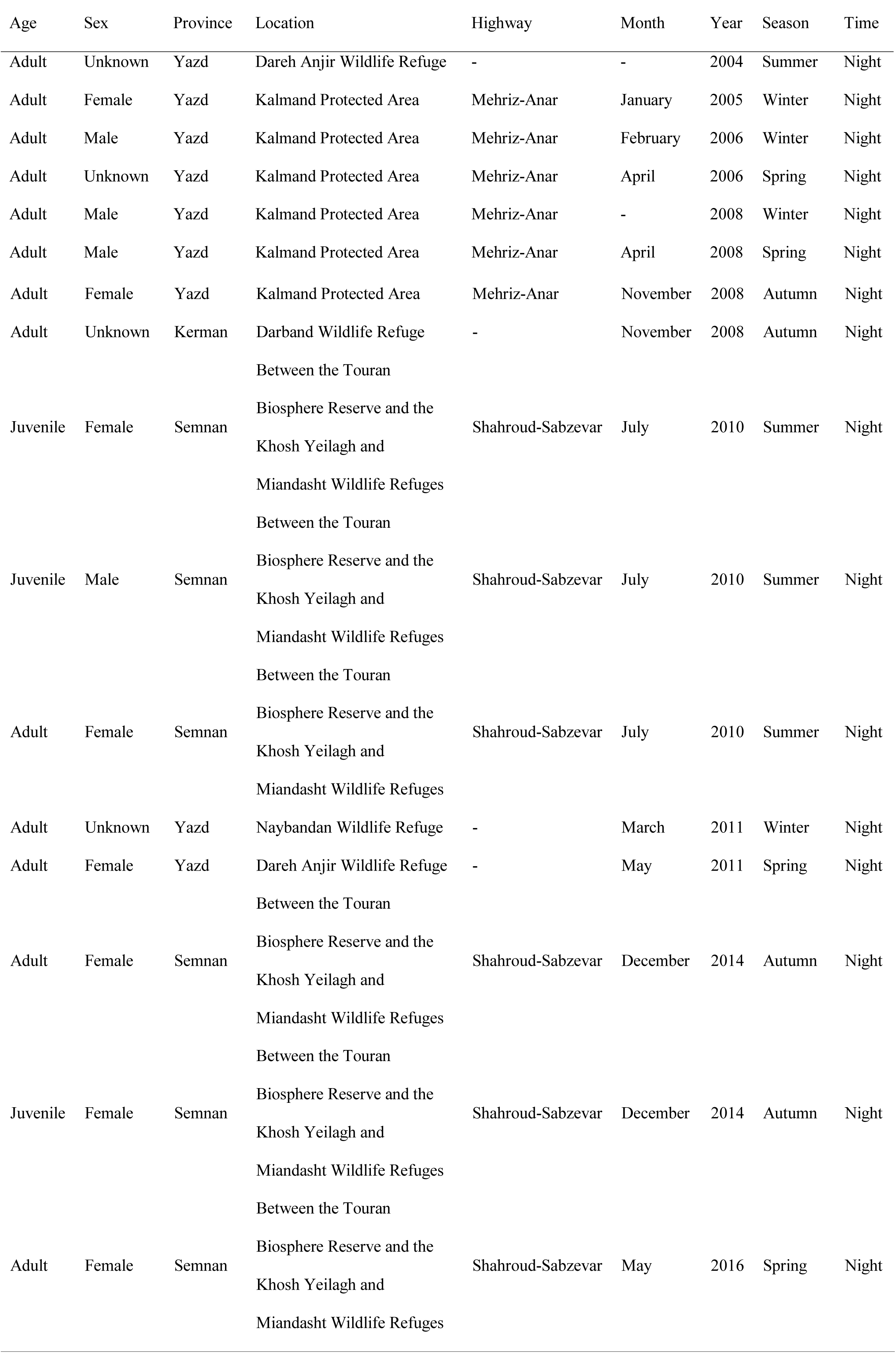
Details of Asiatic cheetah road fatalities recorded in Iran between 2004 and 2016

**Fig. 2.**
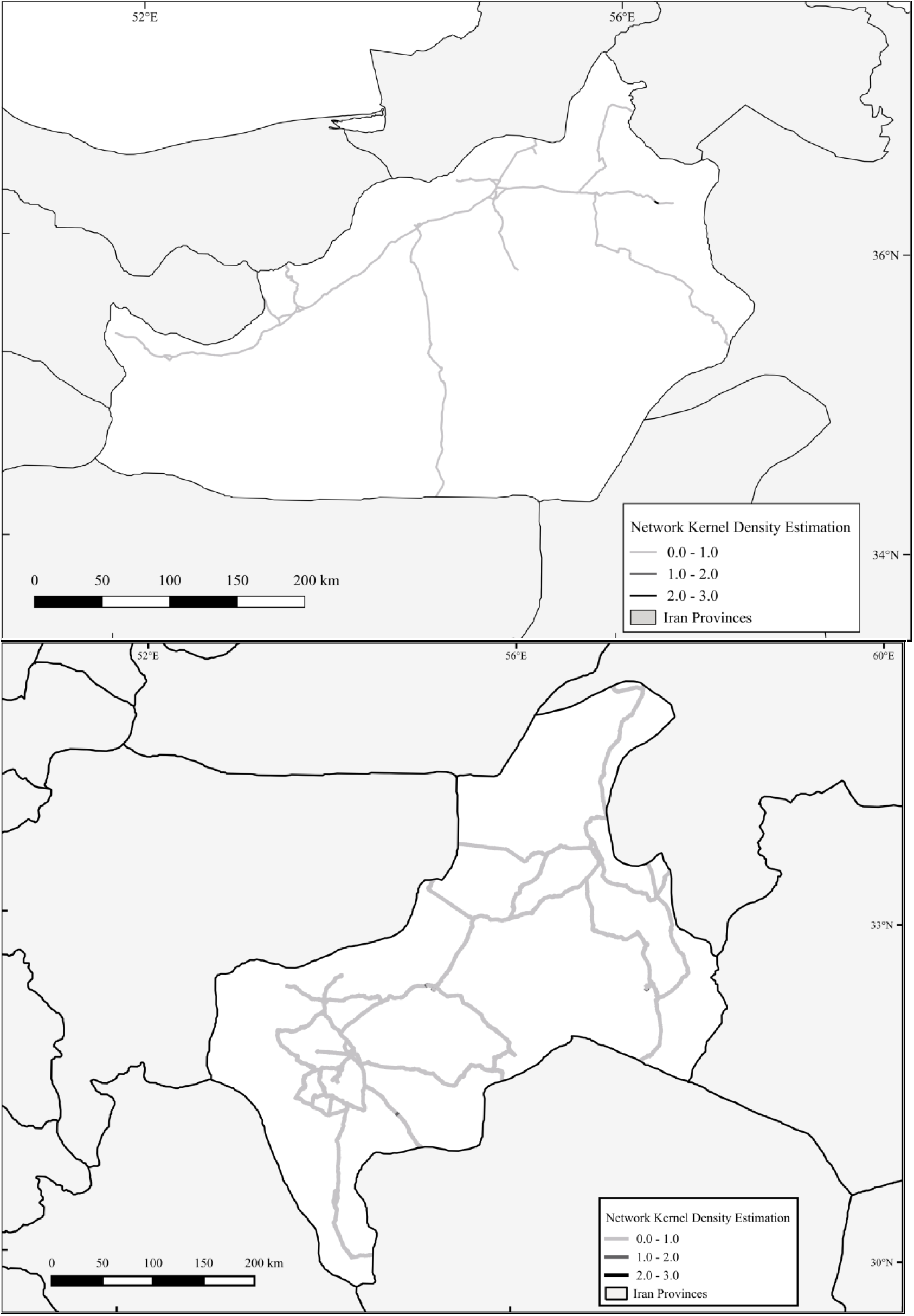
Map showing the locations of the cheetah roadkill hotspots identified using network kernel density estimation in a) Semnan Province, and b) Yazd Province

All fatalities occurred at night and mostly along two major highways that run through or between protected areas: the Shahroud-Sabzevar Highway in Semnan Province, which passes between the Touran Biosphere Reserve and the Khosh Yeilagh and Miandasht Wildlife Refuges, and the Mehriz-Anar Expressway, which passes through the Kalmand Protected Area in Yazd Province (Fig. 1).

Of the 16 collisions three involved juveniles and the rest involved adult animals (six adult females, three adult males, and four adults of unidentified sex). There was no seasonal pattern, with collisions occurring in all seasons and most months of the year, and there were also no clear longitudinal trends in the number of collisions recorded as the study progressed (Table 1).

## Discussion

Hotspots for Asiatic cheetah mortalities due to wildlife-vehicle collisions tended to be located in close proximity to protected areas in which the species occurs. Most cheetah roadkill events were recorded on two major highways (the Shahroud-Sabzevar Highway and the Mehriz-Anar Expressway), possibly because these two roads have a high volume of traffic and were upgraded to dual carriageways approximately 20 years ago. Population growth and urbanization increase the need for the road network expansion, and road construction in ecologically sensitive habitats such as protected areas and national parks has been increasing in Iran during the past decades (Monavari and Mirsaeed 2008; Makki et al. 2013; Mohammadi and Kaboli 2016). Similar trends have also been observed following the upgrade of Cradle Mountain Road in Tasmania, Australia, which led to the local extinction of eastern quolls (*Dasyurus viverrinus*) and to the severe reduction of populations of the Tasmanian devil (*Sarcophilus harnsii*) (Jones 2000).

The high level of road mortality of Asiatic cheetahs is consistent with previous research, which estimated that collisions with vehicles represented the greatest source of cheetah mortality in Iran (42% of recorded mortalities; Ahmadi et al. 2017). Road mortality appears to be much higher for Asiatic cheetahs than for many other large carnivores, and has put them at risk of extinction (Ghoddousi et al. 2017). For example collisions with vehicles were responsible for only 2% of the recorded deaths of Amur tigers (*Panthera tigris altaica*) in Primorski Krai Province, Russia; and 6-26% of leopard (*Panthera pardus*) deaths in Southern Africa (Goodrich et al. 2008; Swanepoel et al. 2015).

Cheetahs may be more susceptible to road mortality than other large carnivores as they have very large home ranges, so they encounter roads more frequently than species with more localized home ranges (Durant et al. 2017). The high level of Asiatic cheetah mortality on Iran’s roads may also be linked to road use by cheetahs. In response to threats such as persecution, poaching, habitat loss, and reduced prey availability, Asiatic cheetahs may alter their use of space by dispersing long distances from their natal ranges (Palomares and Caro 1999; Marker et al. 2008; Farhadinia et al. 2017). Thus, it is possible that cheetahs are killed in road collisions as they utilize Iran’s roads as migration corridors to travel between protected areas (Kerley et al. 2002).

All of the road mortalities occurred at night, even though throughout most of their range cheetahs are normally most active during daylight hours (Broekhuis et al. 2014). Cheetahs may, however, be most active nocturnally in desert environments or areas in which they are persecuted, in order to avoid heat exhaustion and conflict with humans (Marnewick et al. 2006; Belbachir et al. 2015). Crossing roads at night could also be a strategy to minimize risk, as traffic volume is typically lower during the night (Clevenger et al. 2003).

We briefly review the options available to mitigate cheetah fatalities on Iran’s roads, and provide practical recommendations on how this could be best achieved.

### 1. Wildlife crossing structures

Wildlife crossing structures can be very effective at reducing road mortality, especially when used in conjunction with fencing (Grilo et al. 2015). At present there are no wildlife crossing structures in Iran, but we recommend that installing wildlife crossing structures integrated with fencing should become the focus of the road fatality mitigation efforts for the Asiatic cheetah. Care should be taken to design the wildlife crossing structures to maximize their use by cheetahs (Smith et al. 2015). Larger structures, for example, tend to have higher rates of use by carnivores than smaller ones (Kusak et al. 2009). There are many different types of wildlife crossing structures, but of the designs available to them, pumas (*Puma concolor*) appeared to have a preference for open-span bridge underpasses over other types (Gloyne and Clevenger 2001). The location of wildlife crossing structures may, however, be more important than their design (Andis et al. 2017). Crossing structures should be constructed where they would have maximum impact, such as at the hotspots identified, or in areas frequently crossed by cheetahs, or along roads that bisect areas with high cheetah utilization, density or occupancy (Smith et al. 2015), following best practice guidelines (Trocmé 2015). Crossing structures are commonly used by large carnivores (Grilo et al. 2008, Andis et al. 2017) so it is reasonable to expect that Asiatic cheetahs will also use them, although confirming this would require monitoring. While wildlife crossing structures are successful at reducing road mortality, they do not keep animals off the road: to achieve this fencing is also required.

### 2. Fencing

Fencing can be very effective at keeping animals away from roads and funnelling them towards safe areas to cross such as wildlife crossing structures, thus reducing the rate of roadkill (Seiler 2005). Fencing can also, however, fragment habitats and reduce gene flow (Riley et al. 2006). This is particularly problematic for small populations of rare species, and could result in reduced genetic diversity due to inbreeding depression. This has the potential to increase vulnerability to disease, for example, which could contribute to population declines and further increase extinction risk (Benson et al. 2016). As a result, it is imperative that fencing is used in conjunction with wildlife crossing structures in order to maintain connectivity (van der Ree et al. 2015). It is also important to incorporate structures such as jump-outs into fencing designs to allow animals that breach the barrier to return to safety (Huijser et al. 2009). Iran’s Ministry of Roads and Urban Development is planning on fencing 12 km of the Shahroud-Sabzevar Highway over the next two years to secure the most dangerous section of the highway (Abdollahpour 2017). While we welcome this development, we recommend that fencing should be integrated with wildlife crossing structures to ensure landscape connectivity. Consideration of the optimal fencing design for the target species is also important. Fencing designs aimed at reducing roadkill rates for large felids should be used where possible, such as this design used for the Florida panther (*Felis concolor coryi*) (Foster and Humphrey 1995).

### 3. Animal detection systems

Animal detection systems can display a warning message to motorists when target animals are detected using electronic sensors, ensuring that warnings are very specific in time and space. Although these systems could potentially be very effective at reducing wildlife-vehicle collisions involving large animals (Huijser et al. 2015), they also have important drawbacks (Huijser et al. 2015): 1) the rate of reduction in wildlife-vehicle collisions for animal detection systems was 33-97%, compared to 80-100% for fencing integrated with wildlife crossing structures; 2) their implementation requires greater investment in the long term; 3) they are still considered experimental rather than a proven mitigation measure; and 4) they do not reduce the barrier effect of roads and traffic to wildlife. Detection systems are only able to detect larger mammals, and it seems likely that the system would be effective for cheetahs (M. Boyce, pers. comm.; M. Huijser, pers. comm.). Therefore this could be a measure worth considering for the Asiatic cheetah in the future, but it requires further investigation.

### 4. Improving lighting

Enhancing road lighting has sometimes been employed to reduce wildlife road fatalities, as better visibility could enhance the ability of drivers to avoid collisions with wildlife. The effectiveness of this, however, has not yet been clearly proven (Mastro et al. 2008). Furthermore, road lights have significant negative effects on the environment (Blackwell et al. 2015) and do not keep animals off roads, so we would recommend them only if fencing with wildlife crossing structures cannot be installed.

### 5. Enhance warning signage

To date the main approach taken to reduce road mortality for the Asiatic cheetah has been centred on the use of standard and enhanced wildlife warning signs. The authorities installed 16 warning signs on the Shahroud-Sabzevar Highway in Iran in 2013. Such signs are one of the most frequently used measures to mitigate wildlife road fatalities due to their low cost (Huijser et al. 2015). There is little evidence, however, that they are effective at reducing roadkill (Bullock et al. 2011), and it appears that the cheetah warning signs installed in Iran did little to reduce the rate at which cheetahs are killed on the roads. Wildlife warning signs that are spatially- and temporally-specific are more likely to be effective at reducing roadkills (Huijser et al. 2015). Iran’s Ministry of Roads and Urban Development will install 25 wildlife warning signs with flashing lights before and after the 12 km fences on the Shahroud-Sabzevar Highway to reduce road mortalities involving cheetahs (Abdollahpour 2017). Flashing lights, however, cannot be considered an effective measure unless very low vehicle speeds (i.e. less than 50-60 km/h at night) can be achieved (M. Huijser, pers. comm.). We therefore do not recommend installing wildlife warning signs with flashing lights since they are highly ineffective at reducing wildlife-vehicle collisions involving cheetahs. We only recommend road signs that are specific in time and space (M. Huijser, pers. comm.) if fencing with wildlife crossing structures cannot be installed.

### 6. Reducing driver speed

High vehicle speed is an important factor contributing to road mortality (Ramp et al. 2006). The average speed of vehicles passing through Shahroud-Sabzevar Highway and Mehriz-Anar Expressway exceeds 120 km/h at night and is 110 km/h during the day. Reducing vehicle speed, however, is impractical on large, high-speed roads (Smith et al. 2015), such as those on which we have identified roadkill hotspots. Although reducing driver speed at hotspots could help to reduce cheetah mortality rates, introducing lower speed limits and implementing traffic calming measures such as speed bumps are unlikely to be acceptable solutions (M. Huijser, pers. comm.). Iran's Ministry of Roads and Urban Development is planning on implementing traffic calming measures before and after the 12 km fences on the Shahroud-Sabzevar Highway to reduce road fatalities involving cheetahs (Abdollahpour 2017). However, we do not recommend this measure since it could be largely ineffective.

Based on this brief review of potential mitigation strategies we recommend a strategic shift in efforts to reduce road mortality of the Asiatic cheetah. The focus of mitigation should move away from wildlife warning signs, and centre on constructing wildlife crossing structures in conjunction with fencing, as these are likely to be much more effective (Andis et al. 2017). It is important to evaluate and monitor the effectiveness of mitigation strategies (Smith and van der Ree 2015). Although there are no studies available on the use of wildlife crossing structures by cheetahs, it is likely that they would use them if available, as has been observed in other large carnivores such as pumas, black bears ( *Ursus americanus*), grizzly bears (*U. arctos*), and wolves (*Canis lupus*) (Gloyne and Clevenger 2001; Ford et al. 2009; Benson et al. 2016). If monitoring demonstrated that the initial wildlife crossing structures and fencing were used as anticipated by cheetahs, this scheme could be expanded to a broader scale. Tools such as camera traps or microchip readers could be used to demonstrate their use by multiple individuals, a crucial step in demonstrating their use at a population level (Andis et al. 2017; Marnewick et al. 2008). Camouflaging and securing camera traps to fixed objects may help to reduce theft, which can be common as poor availability in Iran makes them a target for theft (Parchizadeh 2017a, b).

Installing wildlife crossing structures and fencing, and monitoring their effectiveness, will require a much greater level of investment than has previously been available to install warning signs. Building wildlife crossing structures is extremely costly (Popescu 2017), although modifying existing infrastructure can be both successful and cost-effective (Mata et al. 2008). Plans must also be made for maintenance of any infrastructure if it is to be an effective solution in the long-term (van der Ree et al. 2015). One source of funding could be insurance schemes, such as that provided by Dana Insurance Company, which contributes up to USD 14,000 per cheetah mortality caused by road collisions or herder dogs (Mehr News Agency 2016). Additional contributions from the Iran's Ministry of Roads and Urban Development would facilitate the implementation of these measures, and funding of mitigation measures should become a standard component in road development budgets (Collinson 2013).

Environmental issues such as strategies to reduce wildlife mortality should be considered during pre-construction planning of a road project, and should be implemented during the early construction phase by installing permanent wildlife crossing structures and fencing (Weller 2015). However, these stages are generally overlooked in Iran, and as a result the Iran’s Ministry of Roads and Urban Development will spend about USD 530,000 to retrofit fences, 25 wildlife warning signs with flashing lights, and traffic calming measures on approximately 14 to 15 km of the Shahroud-Sabzevar Highway over the next two years, which may not be effective at reducing cheetah road fatalities. Additional support from non-profit organizations, or from private organizations operating road concessions could also prove useful, as has been the case in South Africa (Williams et al. unpublished data). Public-private partnerships can also be a key to supporting such projects, such as the current campaign to construct a wildlife crossing to protect cougars in the Santa Monica mountains in Los Angeles, USA (Popescu 2017). Such schemes in Iran could also potentially benefit from innovative funding models that have recently been gaining interest, such as the species royalty system or World Heritage Species concept (Wrangham et al. 2008; Good et al. 2017).

In addition to funding, institutional support is another important component necessary for effective of carnivore conservation (Treves et al. 2017). Worryingly, Iran’s Department of the Environment recently declared that cheetah conservation was no longer their priority, and the United Nations Development Program announced that it is withdrawing from the crucial Conservation of the Asiatic Cheetah Project (Parchizadeh and Williams 2017). This could signal lack of political will and a fall in both domestic and international support for Asiatic cheetah conservation, which could have disastrous consequences for the species. It is important that the Department of the Environment and the international community reaffirm their commitments to Asiatic cheetah conservation if their extinction is to be avoided.

In conclusion, the roads of Iran pose the most serious threat to the Asiatic cheetah. A total of 16 cheetah mortalities were recorded between 2004 and 2016, which were concentrated on the Shahroud-Sabzevar Highway and the Mehriz-Anar Expressway. The loss of a single Asiatic cheetah from a remaining population of only 43 individuals on Iran’s roads can have huge impacts on this critically endangered sub-species, so mitigation efforts are critical. We therefore recommend targeting roadkill mitigation measures at the hotspots identified, and we advocate a shift in strategy away from using warning signage and towards installing wildlife crossing structures in conjunction with fencing. We hope that taking an evidence-based approach and adopting these measures will go a long way towards reducing the threats posed by roads to the Asiatic cheetah.

